# Xenon plasma focused ion beam lamella fabrication on high-pressure frozen specimens for structural cell biology

**DOI:** 10.1101/2024.06.20.599830

**Authors:** Casper Berger, Helena Watson, James Naismith, Maud Dumoux, Michael Grange

## Abstract

Cryo focused ion beam lamella preparation is a potent tool for *in situ* structural biology, enabling the study of macromolecules in their native cellular environments. However, throughput is currently limited, especially for thicker, more biologically complex samples. We describe how xenon plasma focused ion beam milling can be used for routine bulk milling of thicker, high-pressure frozen samples during lamellae preparation with a high success rate and determine a 4.0 Å structure of the *Escherichia coli* ribosome on these lamellae using sub volume averaging. We determine the effects of increased ion currents on sample integrity during bulk milling of thicker planar samples, also showing that beyond an initial region of damage, no measurable structural damage propagates beyond this. The use of xenon results in substantial structural damage to particles up to 30 nm in depth from the milled surfaces, with detectable damage observed to 45 nm. Ours results outlines how the use of high currents using xenon plasma focused ion beam milling may be integrated into FIB milling regimes for preparing thin lamellae for high-resolution *in situ* structural biology.

## Introduction

Cryo-electron tomography, combined with focused ion beam (FIB) lamella fabrication, has become an effective method to determine how macromolecular structure and biological function are linked in the cell (Berger et al., 2023; Young and Villa, 2023). Advances in automated lamella fabrication (Zachs et al., 2020; Klumpe et al., 2021; Tacke et al., 2021), cryo-electron tomography data collection (Hagen et al., 2017; Eisenstein et al., 2023; Khavnekar et al., 2023) and sub-tomogram averaging (STA) (Tegunov et al., 2021; Zivanov et al., 2022) have enabled structures of large macromolecules present at high cellular concentrations to be determined at pseudo-atomic resolution (Sutton et al., 2020; Hoffmann et al., 2022; Wang et al., 2022; Berger et al., 2023; Xing et al., 2023). Further improvements in quality and throughput of complex lamella sample preparation approaches would benefit *in situ* structural biology by resulting in an increase in the attainable resolution for abundant macromolecules and bringing rarer macromolecules within scope.

The limitations of gallium liquid metal ion sources for FIB lamella fabrication have been previously discussed in the context of material science applications (Howitt, 1984; Prenitzer et al., 2003; Smith et al., 2006). In summary, the limited current density of these sources significantly limits the rate of bulk milling. This is a significant drawback for production from thicker, more biologically complex samples such as tissues and small organisms (Schiøtz et al., 2023). Plasma sources however remain collimated even at high currents (Smith et al., 2006) enabling the use of higher current regimes (up to 2.5 µA); the effect upon biological samples remains to be determined. Plasma ion sources have been demonstrated as an effective alternative to gallium sources for lamella production from mammalian cells (Berger et al., 2023) and protein crystals (Martynowycz et al., 2023; Parkhurst et al., 2023). Xenon has a higher milling rate on vitreous samples compared to other ion sources (Fu et al., 2008; Berger et al., 2023), which in combination with the range of currents possible for plasma focused ion beams (PFIB) may enable novel high-throughput milling strategies for thicker biological samples (Schiøtz et al., 2023).

FIB milling is known to damage materials due to ion impacts on the milled surfaces and the resulting collision cascade (Howitt, 1984; Prenitzer et al., 2003; Burnett et al., 2016; Liu et al., 2020). For materials science applications, the main determinants of the depth and extent of the damage layer are known to be the composition of the material being milled, the ion source, ion incident angle and ion accelerating voltage (Howitt, 1984; Prenitzer et al., 2003; Burnett et al., 2016). Using *in situ* sub- tomogram averaging and B-factor analysis (Rosenthal and Henderson, 2003), it was demonstrated that 30 kV argon PFIB milling on biological samples results in damage characterised by reduced information content at depths up to 30 to 45 nm from the milling surfaces (Berger et al., 2023). The damage layer for 8 kV and 30 kV gallium FIB milling has been characterised (Lucas and Grigorieff, 2023; Yang et al., 2023; Tuijtel et al., 2024), reporting penetration depths for the damage between 30 and 60 nm, with reduced depth of damage at lower voltage FIB milling.

We present a workflow for the use of xenon PFIB milling to prepare lamellae from ∼25 µm thick high-pressure frozen biological samples with currents up to 60 nA. The use of high-pressure frozen samples has broad potential as this extends the applications that may be harnessed via cryo-ET (Kelley et al., 2022). With this method we determined the structure of the *Escherichia coli* ribosome to a resolution of 4.0 Å. We show that xenon plasma milling using this protocol results in a damage layer which penetrates up to 45 nm from the milling surfaces and describe the effects of the use of a 60 nA probe for bulk milling. Our results demonstrate that xenon plasma milling with currents up to 60 nA can be integrated into milling workflows for high-resolution *in situ* structural biology.

## Results

### Xenon plasma FIB lamella preparation of high-pressure frozen samples

We adapted previously published methods for lamella fabrication of high-pressure frozen samples (Kelley et al., 2022) to plasma FIB milling. Our method enabled simultaneous site preparation, with benefits in throughput and the removal of ice contamination (Fig. 1, Table 1). We high-pressure froze *Escherichia coli* in solution, using an EM grid without support film as a spacer between two flat-sided planchettes to obtain a vitreous sample with a thickness of ∼25 µm (the thickness of the grid). Ice contamination was frequently seen on the samples. We therefore added an ice removal step where both sides of the grid are imaged with the PFIB beam at low magnification with currents of 4-60 nA for short periods of time (Fig. 1a-c, Table 1, Supplementary Video 1). To effectively produce lamellae from material that is ∼25 µm thick on a square mesh grid, it is essential to remove material at each side of the intended position effectively. Two trenches per lamella site are first milled 90° relative to the grid from the backside of the grid based on the stage limits; the front of the grid is not accessible at a perpendicular angle with the PFIB of the microscope used here (Fig. 1d, e). Xenon PFIB milling has been demonstrated to give higher milling rates for vitrified samples compared to other ion sources (Fu et al., 2008; Berger et al., 2023). We used xenon with currents up to 60 nA for trench milling (compared to 15 nA described by Kelley et al. (Kelley et al., 2022). Next, material is removed below each milling site from the front of the grid at increasingly shallower angles towards the grid surface (45°, 28° and 20° relative to the grid) (Fig. 1f). This leads to progressive removal of the material underneath while not being constrained by the grid bar or the material at the front. To further accelerate trench preparation, we used a routine that utilised a low magnification in the ion beam view to simultaneously prepare multiple trenches within the field of view (Fig. 1e,f). Afterwards, rough, medium and fine automated milling steps were performed using progressively lower currents (Fig. 1g-k and n-q), followed by manual notch pattern milling for stress relief (Fig. 1l,r) and automated final lamellae polishing (Fig. 1m,s).

**Figure 1.**
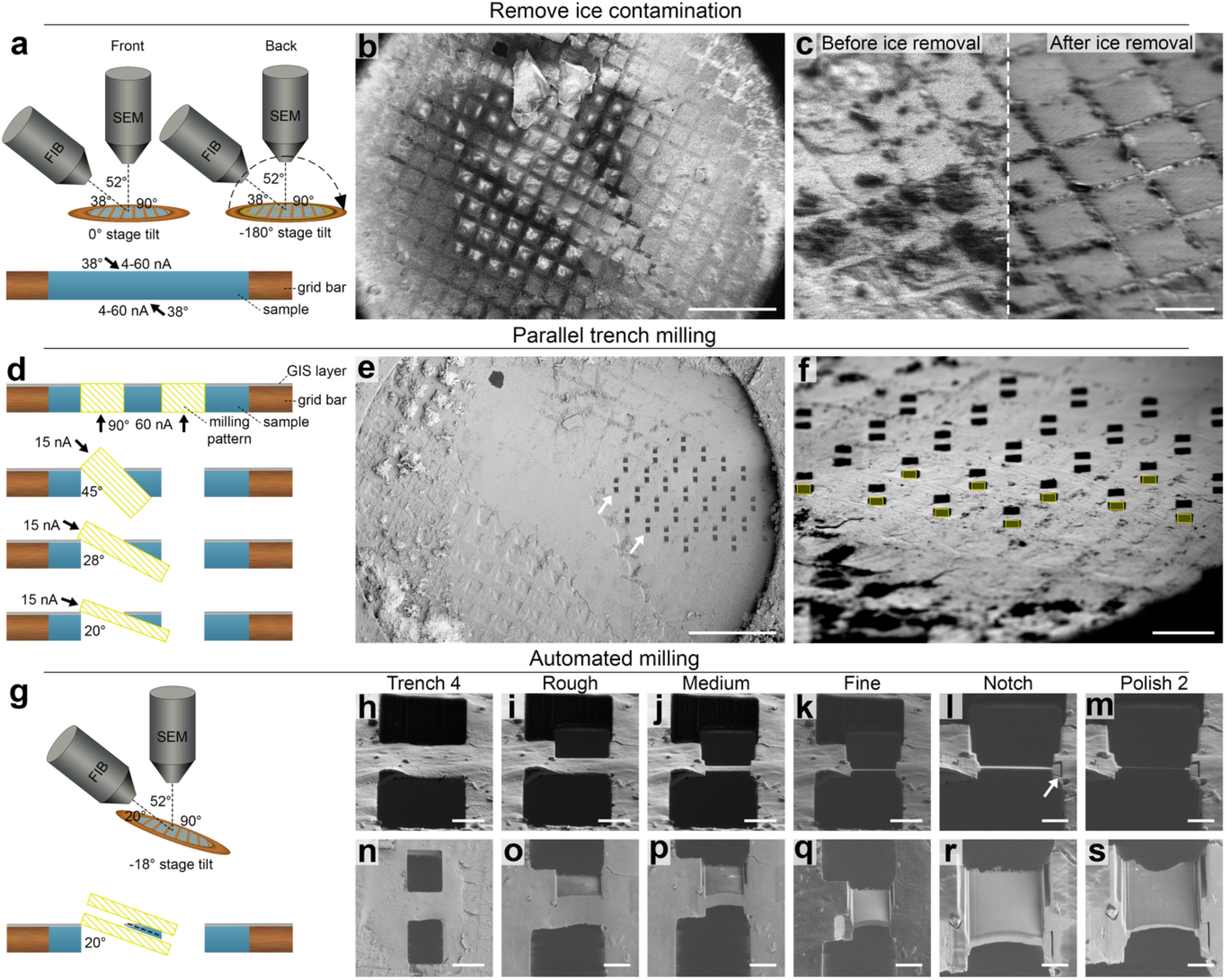
Overview of xenon plasma FIB milling workflow for high-pressure frozen waffle samples of *E. coli*. (a) Orientation of the grid relative to the PFIB and scanning electron microscope (SEM) during ice removal using xenon beam on an Arctis PFIB/SEM from the front (left) and back (right) of the grid. Bottom: schematic orientation of a grid square and the angle and currents used to remove the ice contamination. **(b)** SEM grid overview before ice removal. Scalebar: 500 µm. **(c)** PFIB images before (top left) and after (bottom right) ice removal by PFIB imaging. Scalebar: 100 µm. **(d)** Schematic overview of milling pattern placement (yellow striped rectangles) during trench milling steps. **(e)** SEM overview of the grid shown in **b**, after ice removal, GIS deposition and trench milling steps (example milled sites marked with white arrows). Using a low magnification ion beam, all trenches are milled at the same time. Scalebar: 500 µm. **(f)** PFIB image of pattern placement for the 4^th^ trench milling step. Milling patterns (yellow striped rectangles) are concurrently placed and milled in in horizontal rows, as tilting the grid varies the distance to the PFIB, resulting in a vertical focus gradient. Scalebar: 200 µm. **(g)** (top) Schematic overview of the grid orientation during automated milling relative to the PFIB and SEM beam. (bottom) Schematic overview of milling pattern placement relative to the forming lamella (black dashed line) during automated lamella preparation. **(h-s)** PFIB (top row) and SEM (bottom row) images of an example lamella after the milling step indicated at the top. After the 4^th^ trench milling step (h, n), automated lamella preparation for rough, medium and fine milling (i-k and o-q) at 4 nA, 1 nA and 0.3 nA respectively. After manually preparing notches at 0.1 nA (white arrow) for stress-relief (I, j), two automated milling steps are performed at 30 pA (not shown) and 10 pA (m and s). Scalebars: h, I, j, k: 10 µm. l, m: 5 µm. n: 25 µm. o, p, q: 10 µm. r, s: 5 µm.

**Table 1.** Milling parameters for xenon PFIB milling of high-pressure frozen samples

| Step | FIB angle relative to grid | Pattern width | Pattern height/overlap (%) | Distance between patterns | Pattern type | Milling time* | Drift correction interval | Ion beam current |
| --- | --- | --- | --- | --- | --- | --- | --- | --- |
| Ice cleaning front | 52 | 980 $\mu\text{m}$ by 653 $\mu\text{m}$ | - | - | imaging | ~30-90s | - | 4, 15 and/or 60 nA |
| Ice cleaning back | 90 | 980 $\mu\text{m}$ by 653 $\mu\text{m}$ | - | - | imaging | ~30-90s | - | 4, 15 and/or 60 nA |
| Trench milling 1 | 90° | 25 $\mu\text{m}$ | 30 $\mu\text{m}$ | 25 $\mu\text{m}$ | Cleaning cross section | 3m:00s | - | 60 nA |
| Trench milling 2 to 4 | 45°, 28° and 20° | 22 $\mu\text{m}$ | 18 to 22 $\mu\text{m}$ | - | Rectangular milling pattern | 1m:48s to 2m:12s | - | 15 nA |
| Rough milling | 20° | 20 $\mu\text{m}$** (top)<br>19 $\mu\text{m}$ (bottom) | 8 $\mu\text{m}$ | 6.12 $\mu\text{m}$ | Rectangular milling pattern | 7m:45s | 500 s | 4 nA |
| Medium milling | 20° | 17.3 $\mu\text{m}$** (top)<br>17 $\mu\text{m}$ (bottom) | 250% | 1.72 $\mu\text{m}$ | Rectangular milling pattern | 7m:45s | 500 s | 1 nA |
| Fine milling | 20° | 16.7 $\mu\text{m}$** (top)<br>16.2 $\mu\text{m}$ (bottom) | 240% | 820 nm | Rectangular milling pattern | 7m:24s | 60 s | 0.3 nA |
| Notch | 20° | - | - | - | Line pattern | 50 s | - | 0.1 nA |
| Polishing 1 | 20° | 16 $\mu\text{m}$** | 200% | 260 nm | Rectangular milling pattern | 10m:15s | 60 s | 30 pA |
| Polishing 2 | 20° | 16 $\mu\text{m}$** | 200% | 120 nm | Rectangular milling pattern | 7m:52s | 30 s | 10 pA |

\*For all patterns from one lamella site, but for ice cleaning per grid.
\*\*Typical width of final lamella is 16 $\mu\text{m}$ but can be adapted depending on the lamella site.

### Xenon plasma milling of high-pressure frozen samples results in high-quality lamella for STA

Applying our adapted method to high-pressure frozen *E. coli* cells, 24 sites were prepared, of which 19 were used for automated lamella preparation at 30 kV, and one for manually testing the polishing step (Fig. 2a). 15 of the 19 automatically prepared lamellae were considered suitable for tilt-series acquisition (Supplementary Figure 1), and 286 tilt-series were acquired. We measured the variability in local lamella thickness in each tomogram considered suitable for subsequent STA-based damage analysis (see Materials and Methods) and found a mean local thickness of 200 ± 33.5 nm (Fig. 2b to f, Supplementary Video 2), with lower thickness values at the front compared to the back of the lamellae (Supplementary Figure 2).

**Figure 2.**
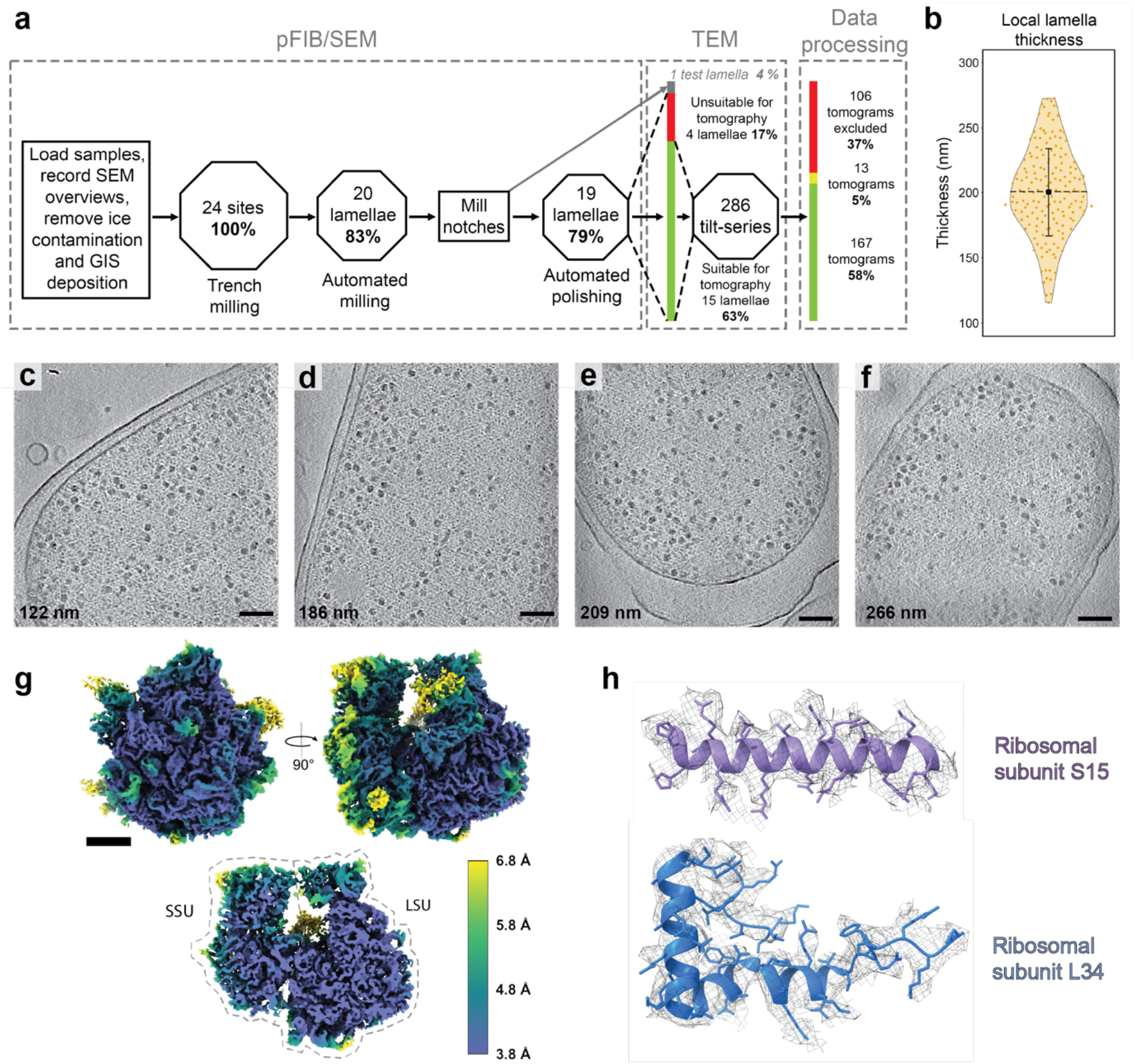
Overview of lamella quality and *E. coli* ribosome STA. (a) Schematic overview of lamella fabrication workflow showing the number of lamellae that were prepared in each milling step, subsequent cryo-electron tomography tilt-series acquisition and which data was used for further analysis. Semi-automated and automated PFIB-SEM steps are indicated in octagons, where the size corresponds to the success rate, and manual steps in rectangles of arbitrary size. Trenches were prepared for 24 lamellae sites, of which 20 were used for automated lamellae fabrication to a thickness of ∼1 µm. Stress-relief notches were prepared manually, followed by automated lamellae polishing for 19 sites (and manual polishing for 1 site for testing milling conditions). The sample was then transferred to the transmission electron microscope (TEM), and 286 tilt-series were collected on 15 out of the 19 lamellae in a 3-day session, using multi-shot beam shift acquisition (Eisenstein et al., 2023; Khavnekar et al., 2023). After data pre-processing and tomogram reconstruction, 106 tomograms unsuitable for subsequent damage analysis were excluded (see materials and methods for the criteria), and a further 13 tomograms could not be further processed during STA due to software errors. **(b)** Violin plot for the local lamella thickness distribution, as measured in the 167 tomograms retained for damage analysis (mean SD;200±33.5 nm, *n*=167). The median (200 nm) is indicated with a black square, the standard deviation with error bars, and the mean with a black dashed line. **(c-f)** Examples of reconstructed slices of tomograms, recorded on lamellae with local thickness values of 122, 186, 209 and 266 nm respectively. Scale bars: 100 nm. **g)** Consensus STA density map of surfaces (top) and central slice (bottom) of 70S ribosome, coloured by local resolution. The small ribosomal subunit (SSU) and large ribosomal subunit (LSU) are indicated. Scale bar: 5 nm. **(h)** Representative model fits into the EM density map for the following ribosomal subunit chains: (L-R) SSU subunit S15, LSU subunit L34.

We extracted ribosomes from our acquired tomograms, resulting in identification of 223,823 predicted ribosome particles (see Materials and Methods). Classification and refinement enabled a consensus 70S ribosome structure with a global resolution of 4.0 Å to be determined (Supplementary Figure 3, Supplementary Figure 4), with large regions of the structure reaching a resolution at the Nyquist sampling limit of 3.8 Å (Fig. 2g, Supplementary Video 3). Indicative of the quality of the map, amino acid side chains could be readily identified in the structure from both small and large ribosomal subunits (Fig. 2h). This 4.0 Å *E. coli* ribosome structure demonstrates the first use of higher current plasma regimes (i.e. incorporating 60 nA steps) for lamella preparation for high-resolution *in situ* structural biology.

### Damage to the backside of lamellae

We observed a region without any apparent biological features on the backside of each lamella (Fig. 3a). This area consists of a striated layer (Fig. 3b, Supplementary Videos 4 and 5), followed by an amorphous region approximately 1 μm in length (Fig. 3b, c). The areas directly adjacent to the amorphous area do contain biological features including membrane bilayers and ribosomes (Fig. 3c, d). We did not observe Bragg reflections in tilt-series recorded on the amorphous layer, indicating that it remains vitreous (Supplementary Video 4).

**Figure 3.**
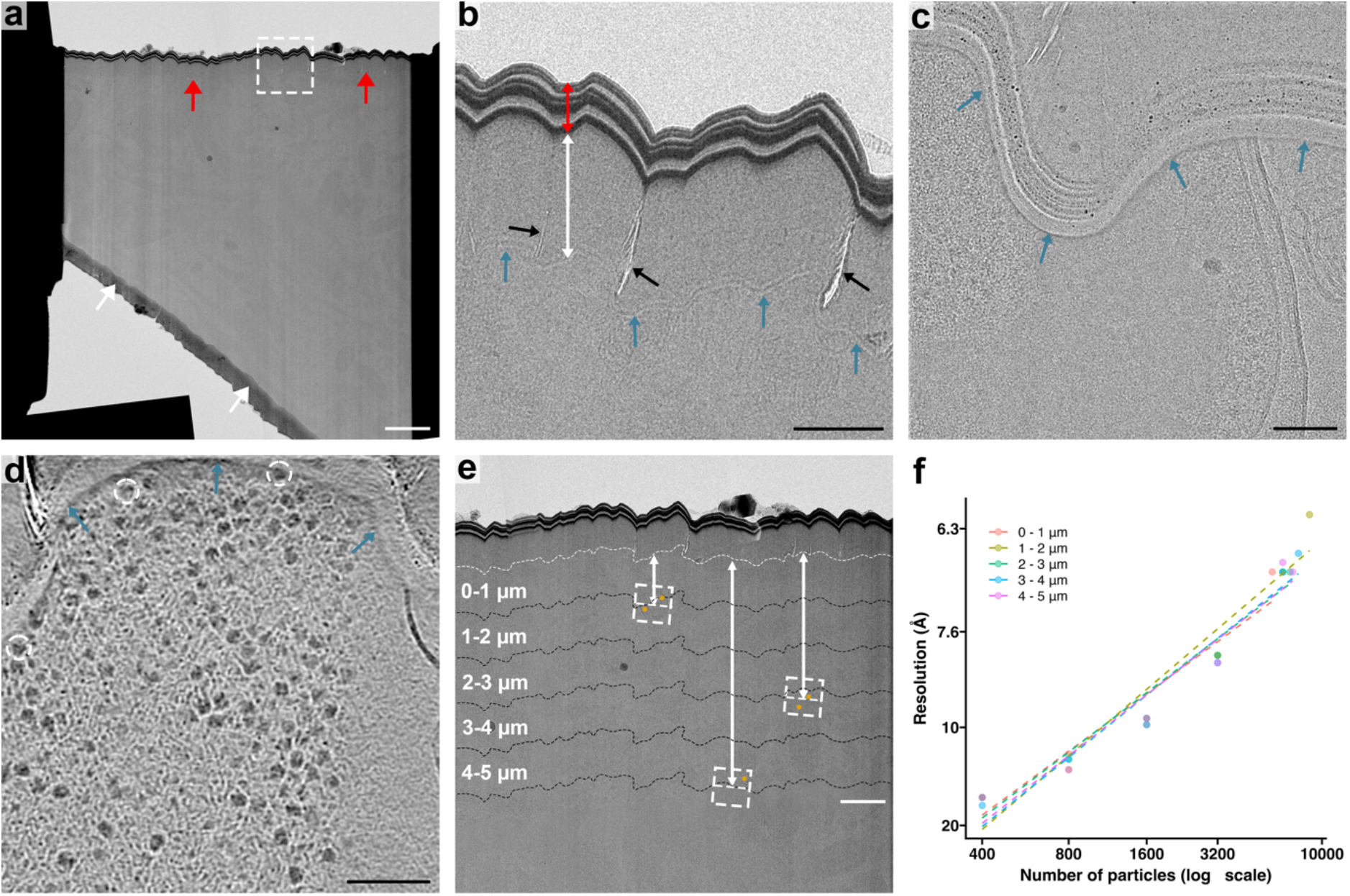
Damage analysis of lamellae backsides. (a) Low-dose TEM overview of a lamella prepared from high-pressure frozen *E. coli* using the protocol described in Figure 1. On the backside of the lamella, a distinct damaged area is visible (red arrows), which is not observed on the front of the lamella, near the GIS layer (white arrows). White dashed square indicates enlarged area shown in panel b. Scalebar: 2 μm **(b)** Enlarged low-dose TEM image of the damaged area on the backside of the lamella, in the area shown in the white dashed square in panel a. A distinct striated pattern of alternating high- and low electron-dense material is visible at the very back of the lamella (red double-headed arrow), with an area without any remaining biological contrast of typically 0.5 to 1.5 μm in length (white double-headed arrow), which often contains small cracks (black arrows). The area without biological contrast is delineated by a striated pattern (blue arrows), weaker in contrast than at the very back of the lamella. Scalebar: 500 nm **(c)** Tilt-image of a bacterium in another lamella, where the contrast of the bacterium abruptly disappears at the border of damaged area without biological contrast (blue arrows). Scalebar: 100 nm. **(d)** Tomographic slice of a bacterium on another lamella, near the area without biological contrast (blue arrows), with ribosomes clearly visible directly adjacent to this border (white dashed circles). Scalebar: 100 nm. **(e)** Enlarged TEM low-dose overview of the lamella shown in panel A, to illustrate how ribosome distance to the boundary with the amorphous layer (white dashed line) is determined per particle, to process them in the corresponding distance group for the B-factor analysis. For each tilt-series (white dashed squares), the distance to the boundary is measured (double headed white arrows), which is modified per-particle based on its coordinates. Scalebar: 1 μm. **(f)** B-factor plot for all ribosomes 1 to 5 μm from the amorphous layer on the backside of the lamellae, in 1 μm groups.

We reasoned that the striated and amorphous regions were damaged by the 60 nA PFIB current used during the perpendicular trench milling step (Supplementary Figure 7). To determine whether this damaged region extended further into the lamella we carried out structural studies using B-factor analysis. In electron microscopy, B-factor plots show the relationship between the number of particles used for reconstruction and the resulting resolution, and are sensitive to all factors that influence this, such as optics and sample characteristics (Rosenthal and Henderson, 2003). We determined the B- factors of ribosomes located at different distances from the boundary between the amorphous region and the rest of the lamella. The distance of each ribosome from this boundary was calculated by using the distance of each tilt series acquisition area from the boundary, then correcting per-particle based on the refined coordinates of the ribosomes within each tomogram (Fig. 3e). Particles were then grouped by distance in groups of 1 μm from the boundary, and the B-factor calculated for each group up to a total distance of 5 μm (Fig. 3f, Table 2, Supplementary Figure 5). The B-factors only ranged between 286 and 321 Å^2^ across the groups, and no correlation was observed (r = -0.3 p-value = 0.68) between B-factors and distance from the boundary between the amorphous layer and the rest of the lamella (Supplementary Figure 5). At the micron scale, we conclude there was no detectable damage propagation beyond the visible amorphous region.

**Table 2.** B-factor analysis of backside damage

| Backside distance group | Number of particles per distance group | Number of tomograms per distance group | Average lamella thickness (nm)* | B-factor ($\text{\AA}^2$) | Global resolution ($\text{\AA}$) |
| --- | --- | --- | --- | --- | --- |
| < 1 $\mu\text{m}$ | 5702 | 18 | 213 ± 29.3 | 321 | 6.8 |
| 1 – 2 $\mu\text{m}$ | 8655 | 26 | 214 ± 27.9 | 286 | 6.2 |
| 2 – 3 $\mu\text{m}$ | 6941 | 21 | 216 ± 26.5 | 319 | 6.8 |
| 3 – 4 $\mu\text{m}$ | 7623 | 26 | 205 ± 29.5 | 304 | 6.6 |
| 4 – 5 $\mu\text{m}$ | 7146 | 25 | 195 ± 24.6 | 310 | 6.8 |

476 \*Average lamella thickness for each particle per distance group.

### Surface damage from milling with xenon propagates to a similar depth to argon

We quantified the depth of ion beam damage propagation from the milling surface into the lamellae using B-factor analysis again. B-factors were determined for ribosomes grouped by distance from the PFIB milling surfaces (Fig. 4a, Table 3). Higher B-factors (Fig. 4b) and lower resolutions for the same number of particles (Fig. 4c) were observed for ribosomes closer to the milling surfaces compared to those located deeper into the lamellae. As the attainable resolution and B-factor of a given particle subset is also impacted by variability of factors such as local lamella thickness and motion, matched controls for each distance group were used with the same numbers of ribosomes taken at random from the same tomograms but at greater distances from the milling surfaces (Fig. 4a, Supplementary Figure 6). We plotted the ratio of B-factor from a given depth group to its matched control (Fig. 4d), as well as the difference between these groups in global STA resolution for ribosomes based on the B-factors (Fig. 4e). In the first 15 nm from the surface boundary models, large differences in B-factors and resolution between the distance groups and their matched controls are observed, which is likely a combined effect from partially ablating ribosomes (diameter of ∼21 nm) as well as damage from ion impacts and the resulting collision cascade. This damage substantially reduces the information content of particles to a depth of 30 nm from the milling surfaces and has a small but measurable effect to a depth up to around 45 nm.

**Figure 4.**
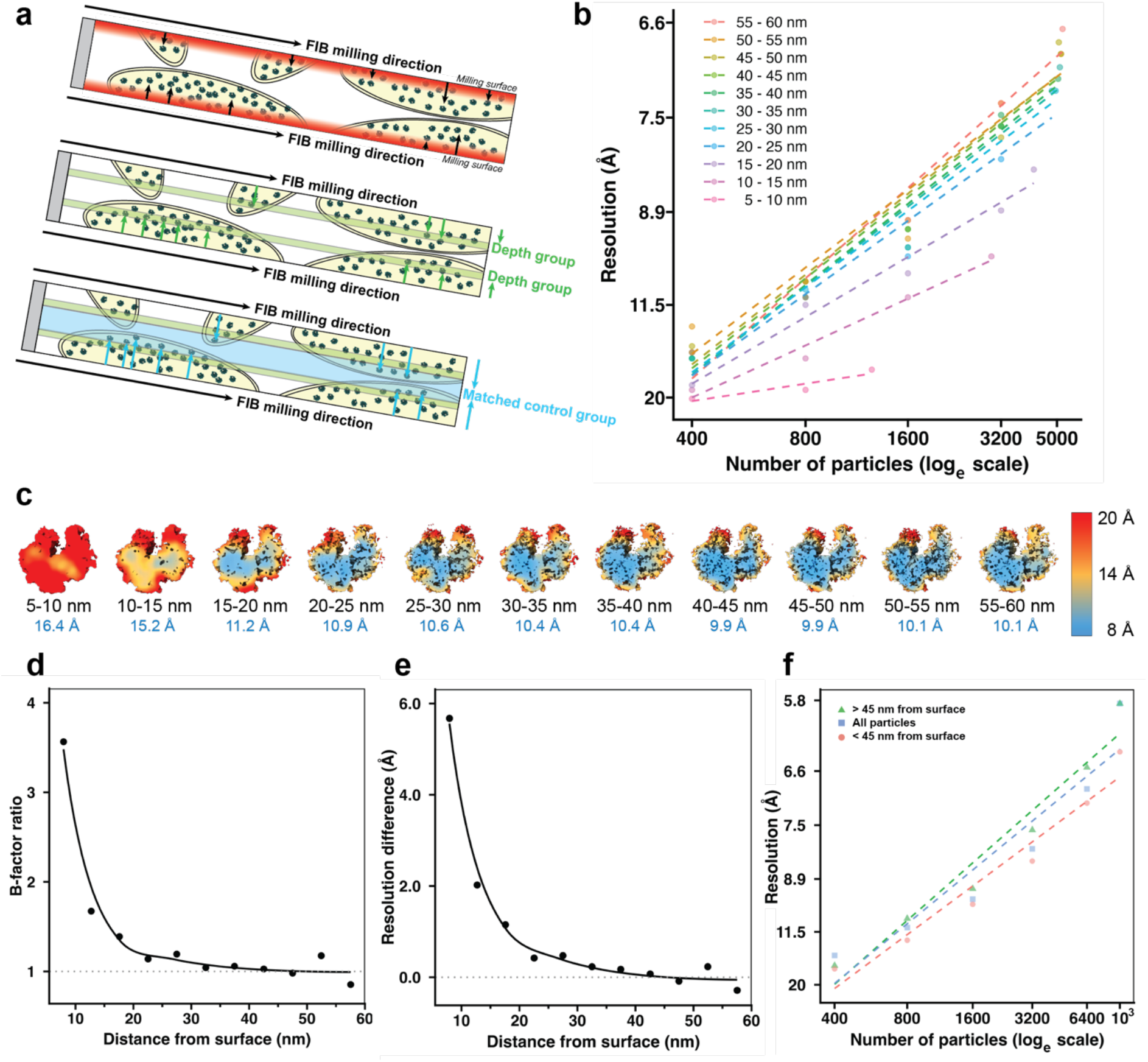
Surface damage layer analysis of xenon plasma milling using STA. (a) Schematic overview of the distance groups and matched controls. For each ribosome, the particle coordinates in the tomogram were used to measure the shortest distance to the annotated milling surface boundary model (top). Ribosomes were then grouped based on their distance to the nearest milling surface in 5 nm bins (e.g. 20 to 25 nm) (middle). To correct for other factors that affect STA resolution (e.g. local lamella thickness, tilt-series alignment quality), matched controls were created for each distance group by randomly taking the same number of particles from the same tomograms, but further away from the milling surfaces (bottom). **(b)** B-factor plots for different 5 nm distance groups from the milling surfaces, up to 60 nm. The 0-5 nm group was excluded from B-factor analysis due to an insufficient number of particles. **(c)** Local resolution maps for the ribosome structures obtained during the B- factor analysis in panel b for 800 particles, with the distance group (black) and obtained global resolution (blue) indicated below each map. **(d)** Plot of the ratio of B-factor for each distance group to its corresponding matched control. **(e)** Plot of the extrapolated difference in resolution for 5000 particles between the distance groups and the matched control groups. **(f)** B-factor plots for 10,000 particles randomly selected from the sets of: all particles, particles less than 45 nm from the milling surface, particles greater than 45 nm from the milling surface.

**Table 3.** B-factor analysis of xenon PFIB milling surface damage

| Surface distance groups | Number of particles per distance group | Number of tomograms per distance group | Average lamella thickness (nm)** | B-factor ($\text{\AA}^2$) | Global resolution ($\text{\AA}$) |
| --- | --- | --- | --- | --- | --- |
| All particles | 10,000 | 163 | $198 \pm 31.6$ | 283 | 5.8 |
| $\leq 45$ nm matched control | 10,000 | 163 | $198 \pm 31.6$ | 315 | 6.4 |
| $>45$ nm matched control | 10,000 | 163 | $198 \pm 31.6$ | 266 | 5.8 |
| 0 – 5 nm | 252 | 91 | $186 \pm 30.2$ | * | * |
| 0 – 5 nm matched control | 252 | 91 | $186 \pm 30.2$ | * | * |
| 5 – 10 nm | 1243 | 152 | $188 \pm 30.0$ | 1491 | 15.8 |
| 5 – 10 nm matched control | 1243 | 152 | $188 \pm 30.0$ | 418 | 9.7 |
| 10 – 15 nm | 2967 | 162 | $186 \pm 29.4$ | 533 | 9.9 |
| 10 – 15 nm matched control | 2967 | 162 | $186 \pm 29.4$ | 319 | 7.6 |
| 15 – 20 nm | 4143 | 161 | $189 \pm 29.7$ | 428 | 8.2 |
| 15 – 20 nm matched control | 4143 | 161 | $189 \pm 29.7$ | 308 | 7.1 |
| 20 – 25 nm | 4795 | 163 | $191 \pm 30.4$ | 351 | 7.2 |
| 20 – 25 nm matched control | 4795 | 163 | $191 \pm 30.4$ | 308 | 6.9 |
| 25 – 30 nm | 4946 | 162 | $192 \pm 30.7$ | 337 | 7.2 |
| 25 – 30 nm matched control | 4946 | 162 | $192 \pm 30.7$ | 283 | 6.7 |
| 30 – 35 nm | 5119 | 162 | $192 \pm 31.0$ | 323 | 7.0 |
| 30 – 35 nm matched control | 5119 | 162 | $192 \pm 31.0$ | 309 | 6.7 |
| 35 – 40 nm | 5048 | 161 | $192 \pm 30.9$ | 319 | 7.1 |
| 35 – 40 nm matched control | 5048 | 161 | $192 \pm 30.9$ | 302 | 6.9 |
| 40 – 45 nm | 5163 | 162 | $192 \pm 31.6$ | 319 | 6.9 |
| 40 – 45 nm matched control | 5163 | 162 | $192 \pm 31.6$ | 310 | 6.8 |
| 45 – 50 nm | 5076 | 163 | $193 \pm 32.1$ | 319 | 6.8 |
| 45 – 50 nm matched control | 5076 | 163 | $193 \pm 32.1$ | 325 | 7.2 |
| 50 – 55 nm | 5179 | 161 | $191 \pm 31.1$ | 305 | 6.8 |
| 50 – 55 nm matched control | 5179 | 161 | $191 \pm 31.1$ | 300 | 6.8 |
| 55 – 60 nm | 5300 | 162 | $194 \pm 30.2$ | 309 | 6.8 |
| 55 – 60 nm matched control | 5300 | 162 | $194 \pm 30.2$ | 322 | 6.8 |

\*Too few particles for reliably determining the B-factor
\*\*Average lamella thickness for each particle per distance group

To determine the overall impact of the damage layer on STA resolution, we performed B-factor analyses for three sets of 10,000 randomly selected particles. One set has particles more than 45 nm from the milling surfaces (no PFIB surface damage), the second set has particles less than 45 nm from the milling surfaces (PFIB surface damaged particles only) and the third set has no distance constraint. STA of particles within 45 nm of the PFIB milling surfaces results in lower resolution structures compared to the other two sets (Fig. 4f).

## Discussion

We have adapted previously published methods for milling high-pressure frozen samples (Kelley et al., 2022) with a xenon PFIB source, improving throughput and implemented an ice contamination reduction step. We can routinely prepare ∼20 lamellae suitable for cryo-ET from high-pressure frozen samples in a 24-hour cryo-PFIB/SEM session, providing comparable success rate, throughput, and lamellae thickness to current methods for plasma- and gallium-milled plunge-frozen cells (Zachs et al., 2020; Berger et al., 2023). This is due to the combination of xenon’s higher milling rate compared to other ion beam sources and the use of higher currents than previously reported for lamella preparation. Even higher currents would further accelerate trench preparation, but samples of this thickness remain rate-limited by the final thinning of lamellae to electron transparency. Higher currents may instead primarily benefit thicker samples such as tissues, which require ablation of much greater volumes of material. To demonstrate the quality of the lamellae produced by our approach we determined a 4.0 Å resolution structure of the *E. coli* 70S ribosome.

One aspect to our approach is the reduction of surface ice contamination with the PFIB; this could have potential to damage the sample. However, we believe this to be unlikely as the large field of view (980 μm x 653 μm) means the estimated milling rate is ∼850 times lower than in the 60 nA trench milling step (25 μm x 30 μm area), and surface damage from the ion collision cascade should be restricted to the front 100-300 nm of the lamella length. The reduction of surface ice allows high quality data collection from grids which would otherwise have been abandoned, with consequent reduction in throughput. The ability to “rescue” grids will particularly benefit projects with complex biological sample preparation where loss of a specimen represents significant experimental effort.

Interestingly, our data show that the use of 60 nA led to an ∼2 μm damaged area on the backside of lamellae, which occurs during the perpendicular trench milling step. However, the front side of the lamellae was undamaged. This is due to the positioning of the lamella site (i.e. typically offset from the front trench by several microns) and the shallow milling angles used (Supplementary Figure 7). As a result, the front side of the lamella is formed from sample that has not been exposed to 60 nA, whereas the lamella backside intersects the back trench side wall. We were unable to perform a suitable B-factor analysis of particles at the front of the lamella (due to insufficient particles), however, B-factor analysis of the lamella backside revealed no detectable damage beyond the visibly affected region (Fig. 3f). Consequently, data acquired adjacent to this region are suitable for high resolution STA.

The mechanism behind the damage to the backside remain to be determined and deeper understanding may help to devise approaches to mitigate or avoid backside damage, especially when considering potential use of larger ion beam currents. Interestingly, no Bragg reflections in these areas were seen within tilt-series, thus the lack of biological contrast is not from devitrification but through a separate mechanism. Based on the abrupt loss of biological features in this region, we hypothesise that exposure to the ion beam has caused rearrangements that disrupt the molecules in the sample without the formation of crystalline ice. We speculate that the type of milling pattern used (e.g. rectangular milling pattern or cleaning cross section) may play a role, as this influences the time frame in which regions of sample are exposed to beam energy, as well as affecting sample redeposition. Further experiments will be required to deconvolve these factors.

The high number of ribosome particles present has allowed finer depth sampling than previously used for argon PFIB surface damage analysis (Berger et al., 2023). In this study, there were no observable effects on the final structure of ribosome of depths (from surface) greater than 45 nm (Fig. 4d,e). We did see an effect on structural quality at depths between 30 and 45 nm but it was relatively small. For depths up to 30 nm there was significant reduction in structural quality (Fig. 4d,e). The limit of the depth of damage is comparable to other studies using argon (Berger et al., 2023; Chaillet et al., 2023; Parkhurst et al., 2023) and gallium beams (Lucas and Grigorieff, 2023; Yang et al., 2023; Tuijtel et al., 2024), despite notable differences in methodology, each of which have relative merits. Regardless of the technique used, it is important to correct for variations in factors such as local lamella thickness, motion and charging. In this study, we use matched controls for the different depth groups as an internal control, to correct for these per-tomogram variables. Considering that current studies conclude comparable ranges in damage penetration depth, other ion-specific factors such as milling rate, probe size and curtaining propensity should be primarily considered when choosing an ion source for lamella preparation.

Methods to reduce the impact of the damage layer on STA require further exploration, particularly where the protein of interest is low abundance, or the biological question requires high resolution. Computationally down weighting particles near the milling surfaces has been shown to improve the reconstruction quality for single particle analysis (Yang et al., 2023), and could be implemented in STA processing pipelines. This would require a highly accurate rapid method to automatically annotate the milling surfaces in tomograms, as manual annotation is low throughput. An alternative is to reduce the accelerating voltage of the ion beam in the final polishing steps, which has precedent in materials science (Howitt, 1984; Burnett et al., 2016) and has recently been demonstrated for gallium FIB for biological samples to reduce the penetration depth of surface damage (Yang et al., 2023). A practical consideration is the increased probe size at lower voltages, which makes it more challenging to prepare thin lamellae and increases the curtaining propensity (Yang et al., 2023). Developing reliable low-kV polishing protocols for preparing high-quality lamellae will be critical to benefit from this reduction in surface damage.

In this work we describe a high-throughput methodology for preparing cryo-FIB lamellae from high- pressure frozen samples with xenon plasma with currents up to at least 60 nA. The approach allowed a 4.0 Å resolution *E. coli* 70S ribosome to be determined using STA. We observed that the use of such currents led to the formation of a ∼2 μm damaged area on the backside of the lamellae, though B- factor analysis suggests that no significant damage propagates beyond the regions that are visibly affected. Our STA and B-factor calculations determined that 30 kV xenon PFIB milling results in detectable damage up to a depth of 45 nm from the milling surfaces, comparable to other ion sources. Overall, this work demonstrates the suitability of xenon PFIB milling for high-resolution *in situ* structural biology on high-pressure frozen samples.

## Methods

### Sample preparation

*E. coli* C43 (DE3) transformed with plasmids as for the plasmid immunity assay specified by Hogrel *et al*. (Hogrel et al., 2022) were incubated overnight at 37 °C in LB supplemented with 100 µg/µl ampicillin, 50 µg/µl spectinomycin and 25 µg/µl tetracycline. The starter culture was used to inoculate a culture at an OD600 of 0.10 in antibiotic-supplemented LB, which was grown to an OD600 of 0.6 to recover exponential growth before the addition of 0.2% (w/v) D-lactose and 0.2% (w/v) L-arabinose at 16 °C overnight to induce expression of the plasmid system. The bacteria were then resuspended with glycerol in LB to a 10% (v/v) final concentration of glycerol and concentrated by centrifugation at 1000 x *g* immediately before vitrification.

Bacteria were vitrified by high-pressure freezing in a Leica EM Ice (Leica Microsystems), following an adapted version of the Waffle Method (Kelley et al., 2022). 3 mm planchettes (Aluminium HPF carrier type B, Science Services) were polished with fine grit sandpaper up to a final grit of 10,000 and metal polish before coating to remove the concentric surface patterns. All planchettes were coated with soya lecithin solution (1% w/v dissolved in chloroform, Sigma-Aldrich) to aid planchette separation after freezing. The Waffle grid was assembled in the following order: planchette hat flat side up, a 200 mesh Cu grid (Agar Grids 200 Mesh Copper 3.05mm, Agar Scientific) glow-discharged on both sides, 5-10 µl of bacterial suspension, planchette hat flat side down. Frozen waffles were disassembled under liquid nitrogen. Vitrified grids were clipped into Autogrids with a cut-out notch (Thermo Fisher Scientific) and stored under liquid nitrogen until lamella fabrication.

### Lamella fabrication

Clipped autogrids were loaded onto an Arctis dual-beam FIB/SEM microscope (Thermo Fisher Scientific). Ice contamination was removed from the front and the back of the grid by FIB imaging with xenon plasma at 15 and/or 60 nA at 30 kV, 38° relative to the grid at the lowest possible magnification (212x), until most of the ice was removed. The front side of the grids was sputter coated with platinum for 120 s at 12 kV and 70 nA, followed by 120 s of coating using the gas injection system (GIS) with tri- methyl(methylcyclopentadienyl)platinum(IV), followed by 120 s of sputter coating with platinum using the same conditions as before. From the backside of the grid at -128° stage tilt (where the FIB is 90° relative to the grid), two trenches were milled per lamella site, where all lamella sites were positioned at low magnification for serial milling. Trenches were prepared with a height and width of 30 and 25 µm respectively, using cleaning cross sections at 60 nA, 30 kV. The distance between trenches for a given lamella site was 25 µm. Material below the lamella sites was removed at 45°, 28° and 20° relative to the grid (7°, -10° and -18° stage tilt respectively) using a rectangular pattern with a width of 22 µm and a height between 18 and 22 µm at 15 nA, 30 kV. Patterns were placed to cover the bottom trench and underside of the lamella. Multiple patterns were concurrently placed in horizontal rows for milling.

After trench milling, lamella sites were identified in Maps (Thermo Fisher Scientific) on an SEM image of the milled grid acquired at eucentric height. Sites were then loaded into AutoTEM Cryo 2.4 (Thermo Fisher Scientific) for automated milling with xenon at 30 kV at a milling angle of 20° relative to the grid. For each lamella site, eucentric height was refined automatically before user determination of the desired final lamella position and width. Three rough milling steps were performed at decreasing currents of 4.0 nA, 1.0 nA and 0.3 nA as the milling patterns were placed with decreasing distance from the intended lamella position (table 1). A notch was then milled manually at 0.1 nA, before re-defining the final lamella site in AutoTEM Cryo. Lamellae were then polished with 30 pA and 10 pA with a distance between the patterns of 260 nm and 120 nm respectively.

### Cryo-electron tomography

Data were collected with Tomo5 software (Thermo Fisher scientific) on a Titan Krios G4 (Thermo Fisher Scientific) TEM with a Falcon 4i camera and a Selectris X energy filter. Low-dose tiled overviews were collected at a 20 degree angle to correct for the lamella pre-tilt and tilt series were collected dose-symmetrically (Hagen et al., 2017) in electron counting mode with the following parameters: magnification of 64,000 x (calibrated pixel size of 1.90 Å/px), tilt range of ± 51° with 3° increments, and a total dose of ∼175 e^-^/Å^2^. The target defocus ranged between 3 and 5 μm across tilt series, in 0.5 μm increments. Tomo5 was used to collect 286 tilt-series in EER format using multi-shot beam shift acquisition (Eisenstein et al., 2023; Khavnekar et al., 2023), with 1 to 6 tilt series acquired parallel, with positions placed on or close to the tilt-axis.

### Tilt series processing

Tilt images with low contrast or objects moving into the field of view (e.g. grid bars or ice contamination) were manually excluded using a custom script. Warp (Tegunov et al., 2021) (version 1.0.9) was used for gain correction, CTF estimation, motion correction using a frame group size of 10 and tilt series stack generation for all 286 tilt-series. Tomogram volumes were reconstructed with AreTomo (Zheng et al., 2022) (version 1.2.5) using simultaneous algebraic reconstruction technique (SART) at a binning factor of 8 and flipped with IMOD (Mastronarde and Held, 2017) (version 4.11.1). For filtering the tomograms we used either a low-pass filtered using EMAN2 (Chen et al., 2019) which thresholds the frequencies, CTF deconvolution using Isonet which uses a Wiener-like filter to boost contrast lost due to the contrast transfer function (Tegunov and Cramer, 2019; Liu et al., 2022), or machine learning based missing wedge compensation using Isonet (Liu et al., 2022). An IsoNet model was trained on 13 selected tomograms, using a cube size of 32 and crop size of 128 without any masking, and applied to all the tomograms.

### Ribosome particle picking

For training a crYOLO (Wagner et al., 2019) model for particle picking, ribosome particles were manually picked all the ribosomes every ∼25 slices from 8 tomograms (3,337 total particles). These co- ordinates were used to train four crYOLO models on (1) unfiltered tomograms, (2) low-pass filtered tomograms, (3) CTF deconvolved tomograms (as implemented in IsoNet), and (4) IsoNet filtered tomograms. Predictions of ribosome co-ordinates on all tomograms resulted in higher mean confidence scores using the model trained on IsoNet filtered tomograms (Supplementary Figure 8), therefore these particle position predictions were used for further processing. This gave 224,823 putative ribosome particles picked above the chosen confidence threshold value of 0.45.

### Sub volume averaging

All 224,823 predicted ribosomes were extracted at a binning factor of 4 (7.6 Å/px) with a box size of 64 pixels using Warp and imported into RELION 4.0 (Zivanov et al., 2022). A subset of 5,415 particle volumes from 6 manually selected tomograms was 3D classified (using the average of this subset as an initial reference), and the most ribosome-like class was used as a reference for 3D refinement of the subset to give a ribosome structure with resolution 16.2 Å. This subset-derived structure was used as a reference for 3D refinement in RELION of the whole dataset, followed by two rounds of 3D classification to identify 95,679 ribosomes. These particles were re-extracted in Warp at a binning factor of 2 (3.8 Å/px) and then 3D refined in RELION to give a ribosome structure with resolution of 7.6 Å.

106 out of 286 tomograms were excluded from further processing using the following criteria: 1) the contrast of the tomogram did not allow for a confident annotation of the milling surfaces, 2) ribosomes could not be clearly identified in the tomogram, or 3) the tomogram contained substantial redeposition or other surface contamination. Particles positions were imported into M and an initial round of M was run with no refinement parameters selected to verify that the programme could run successfully. An additional 13 tomograms were excluded during processing in M due to software errors, resulting in a total of 82,377 particles from 167 tomograms being used for the final reconstruction in M. The following parameters were then applied incrementally over six subsequent rounds of refinement: (1) particle poses, (2) 3x3 image warp grid, (3) 3x3x2x10 volume warp grid, (4) 3x3x2x10 volume warp grid, (5) temporal poses = 3, (6) defocus optimisation with grid search in 1^st^ iteration. This resulted in a ribosome structure with a resolution of 4.0 Å, based on a Fourier-shell correlation using the 0.143 criterion. Panels containing density maps and structures were prepared with ChimeraX (Goddard et al., 2007; Meng et al., 2023).

### Lamella backside damage analysis

Lamella overviews were created by stitching the search images recorded by Tomo5 using the FIJI (Schindelin et al., 2012) plugin “Grid/Collection stitching” (Preibisch et al., 2009). The positions where tilt-series were acquired were found using the first tilt-image recorded for each tilt-series, the central position of each search image in the stitched lamella overview and screenshots of the position placement in Tomo5. Using the measure tool in FIJI, the distance to the boundary where the biological contrast disappears on the backside of the lamella were then measured. This value was then modified per-particle by adding the displacement of the particle X coordinate from the centre of the tomogram to this distance, which was added as a separate column to the .star file. Separate .star files were prepared in 1 µm distance (Table 2) groups using Starparser (https://github.com/sami-chaaban/starparser) and subsets of particles were independently refined in RELION 4.0. B-factors (Rosenthal and Henderson, 2003) were determined for these reconstructions using the Relion bfactor_plot script (https://github.com/3dem/relion/blob/master/scripts/bfactor_plot.py).

### FIB surface damage layer analysis

Damage layer analysis was performed as described previously (Berger et al., 2023). Briefly, lamella surfaces were manually annotated every ∼100 slices and interpolated using a custom script (https://github.com/rosalindfranklininstitute/RiboDist). The script determines the shortest distance for each ribosome to the interpolated boundary models and are saved in a STAR file. Separate STAR files were prepared using Starparser for all ribosomes 0 to 60 nm from the boundary models in 5 nm distance groups (Table 3) and as controls, ribosomes randomly selected from the same tomograms that are further away from the boundaries. If insufficient particles were present in a matched control compared to the corresponding distance group, particles were removed at random from the distance group to obtain the same number of particles as the matched control. B-factors were determined as described above. The 0-5 nm distance group was excluded from B-factor analysis, as too few particles were present.

## Data availability

Subtomogram averages of ribosomes obtained in this study will be deposited in the Electron Microscopy Data Bank (EMDB): full reconstruction (EMD-50675), the reconstructions from the 1 µm distance groups from the backside damage (EMD-50699, EMD-50700, EMD-50701, EMD-50702, EMD- 50703), 5-nm distance groups for the PFIB surface damage (EMD-50676, EMD-50678, EMD-50679, EMD-50680, EMD-50681, EMD-50682, EMD-50683, EMD-50684, EMD-50685, EMD-50686, EMD- 50687) and the matched controls (EMD-50688, EMD-50689, EMD-50690, EMD-50691, EMD-50692, EMD-50693, EMD-50694, EMD-50695, EMD-50696, EMD-50697, EMD-50698), and the 10,000 particle sets for determination of damage layer impact (EMD-50704, EMD-50705, EMD-50706).

## Acknowledgements

We thank Thomas Glen and Matthew Case for microscope support, Neville B.-y.Yee for help with the RiboDist software and Sheera Abdulla for wet lab support. We thank Gaëlle Hogrel and Malcolm F. White for providing the plasmids. We would also like to thank Laura Spagnolo and Malcolm F. White for guidance with bacterial work. This work was supported by the Wellcome Trust through the Electrifying Life Science project (220526/Z/20/Z to J.H.N.) and a Wellcome Career Development Award (225902/Z/22/Z to M.G.). The Rosalind Franklin Institute is funded by UK Research and Innovation through the Engineering and Physical Sciences Research Council (EPSRC).

## Author contributions

H.W., C.B, M.D., and M.G. conceived the study. H.W. prepared cryo samples. C.B. and H.W. prepared PFIB lamellae, collected TEM data, performed STA and damage analyses. C. B., H.W., M.D. and M.G. wrote the manuscript and C.B. and H.W. prepared figures. M.D. and M.G. supervised the project. All authors reviewed the manuscript and the data.

---

**Supplementary Figure 1.**
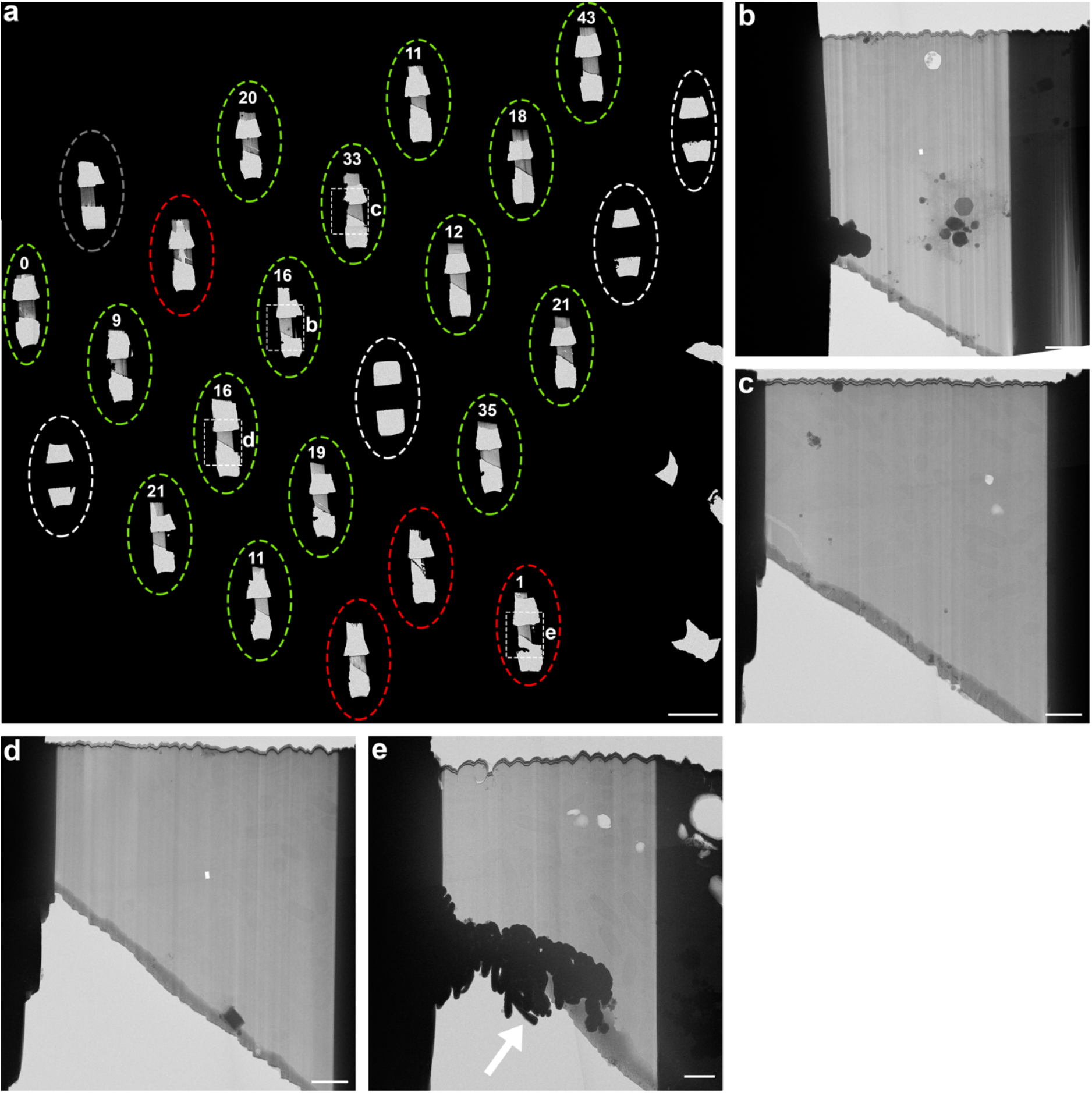
**TEM grid and lamella overviews.** (a) stitched TEM atlas image of a grid containing the 24 lamellae sites indicated in Fig. 2a. Sites that were not further used for automated lamella fabrication are indicated with white dashed ellipses, lamellae considered unsuitable for tilt-series acquisition due to lamella breakage or insufficient removal of material below the lamellae are indicated with red dashed ellipses and tomograms suitable for tilt-series acquisition (clear biological contrast in low-dose TEM overviews on a stable lamella, without objects blocking the beam at high tilts) indicated with green dashed ellipses. One lamella manually polished for test purposes is indicated with a grey dashed ellipse. The number of acquired tilt-series on each lamella is indicated in white above each site, and the white letters on the right site of some sites indicate that low-dose TEM overviews are shown in the respective panel b-e. Scalebar: 50 µm. (b-e) Example low-dose TEM overview of lamellae suitable for tilt-series acquisition (b-d) and unsuitable (e), in this example due to not all the material being removed below the lamella (white arrow). Scalebars: 2 µm.

**Supplementary Figure 2.**
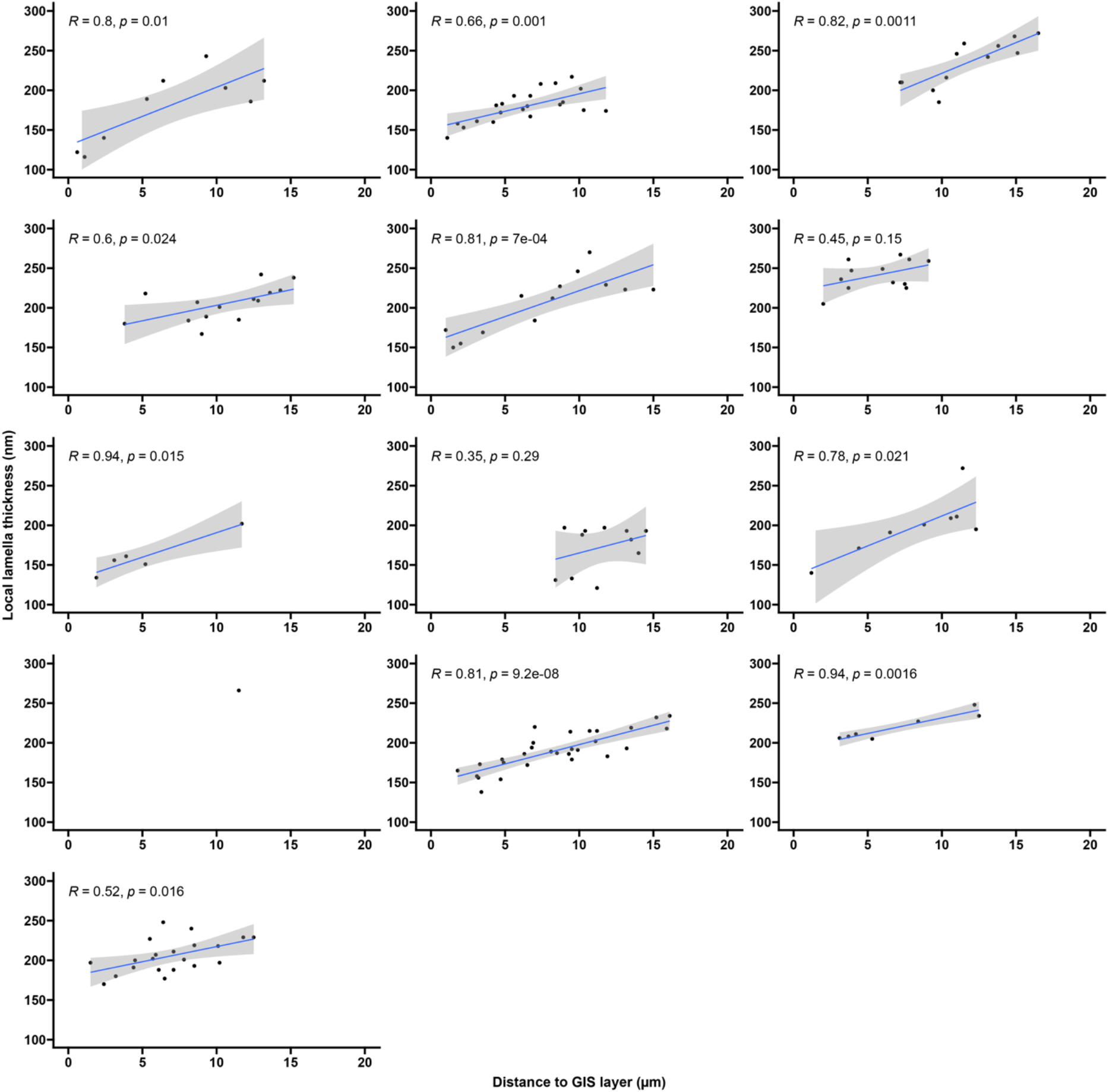
**Plots for local lamella thickness in relation to distance from the front of the lamella.** Scatter plots for local lamella thickness measured in the tomograms used for damage analysis compared to the distance to the front of the lamellae. A linear trend line (blue) is plotted for each lamella with the 0.95 confidence interval (grey) and the Pearson correlation (r) and p-value (p) are shown for each lamella.

**Supplementary Figure 3.**
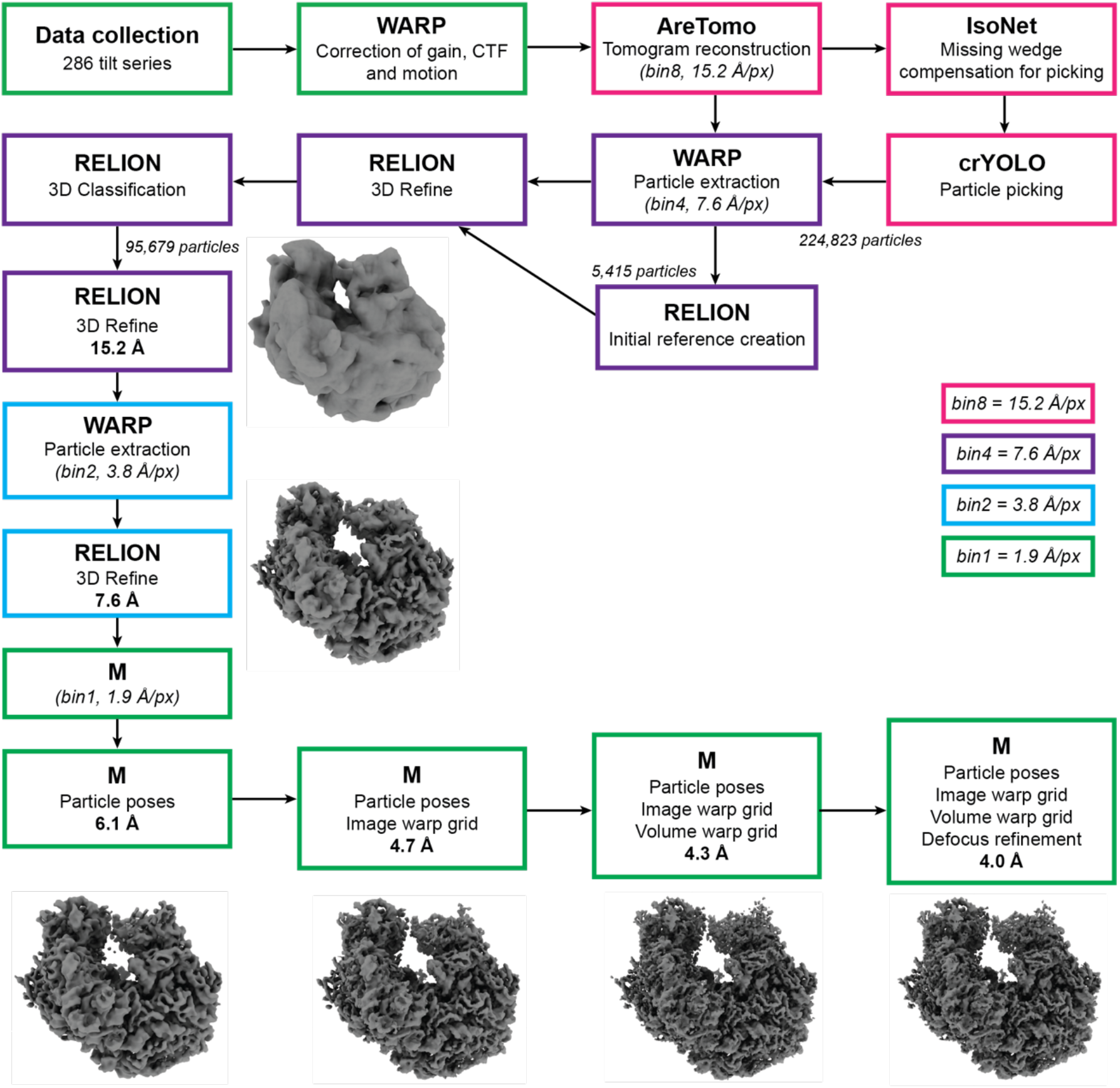
**Overview of STA processing pipeline.** Coloured frames indicate the binning factor used during each processing step and the obtained resolution of the density map of each 3D refinement step is indicated in bold.

**Supplementary Figure 4.**
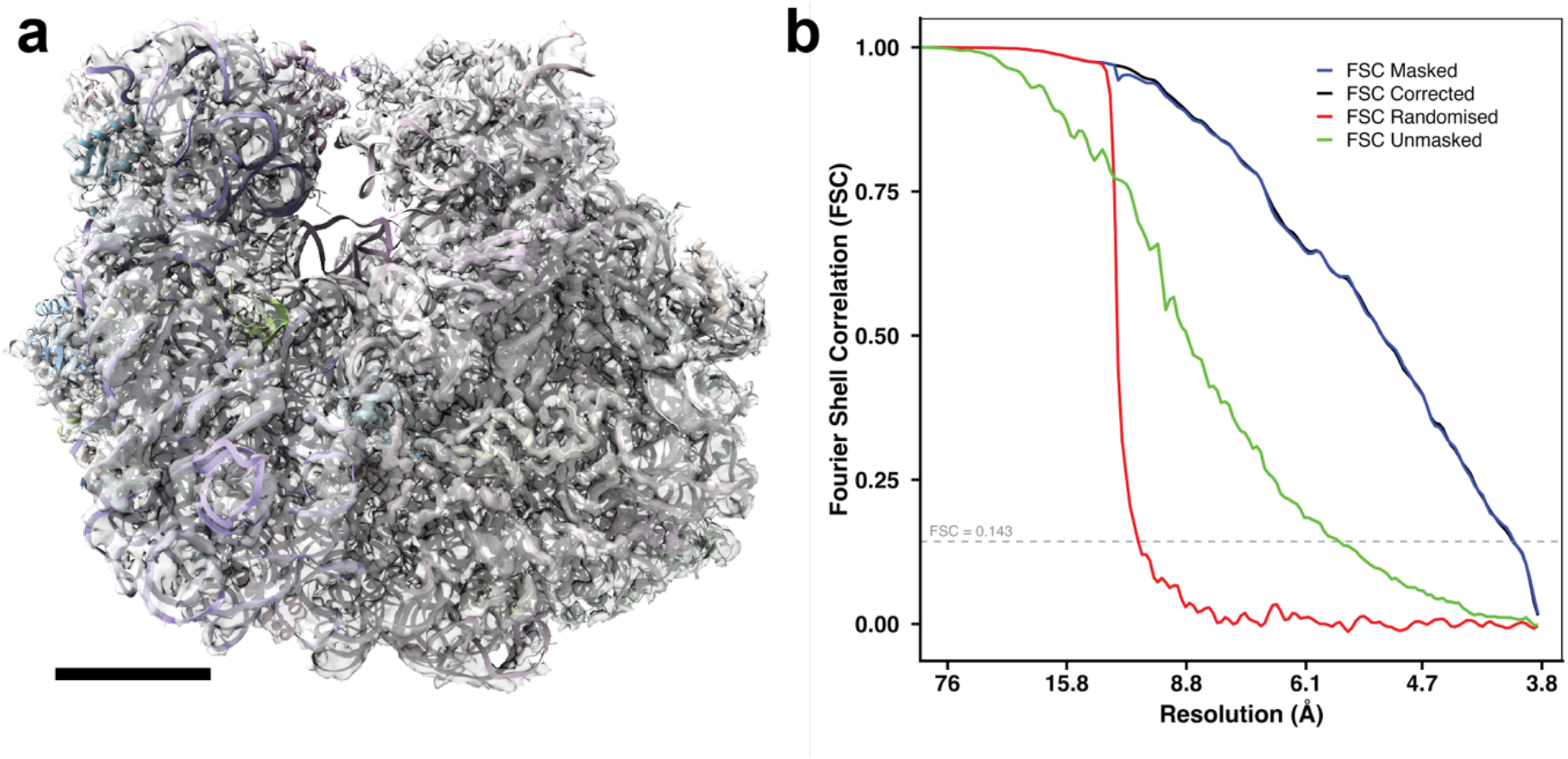
*In situ* STA structure of the *E. coli* 70S ribosome. (a) Consensus STA density map of the 70S ribosome with fitted cryo-EM atomic model (PDB: 6ORE)(Fu et al., 2019). Scale bar: 5 nm **(b)** Fourier Shell Correlation gives masked global resolution of 4.0 Å.

**Supplementary Figure 5.**
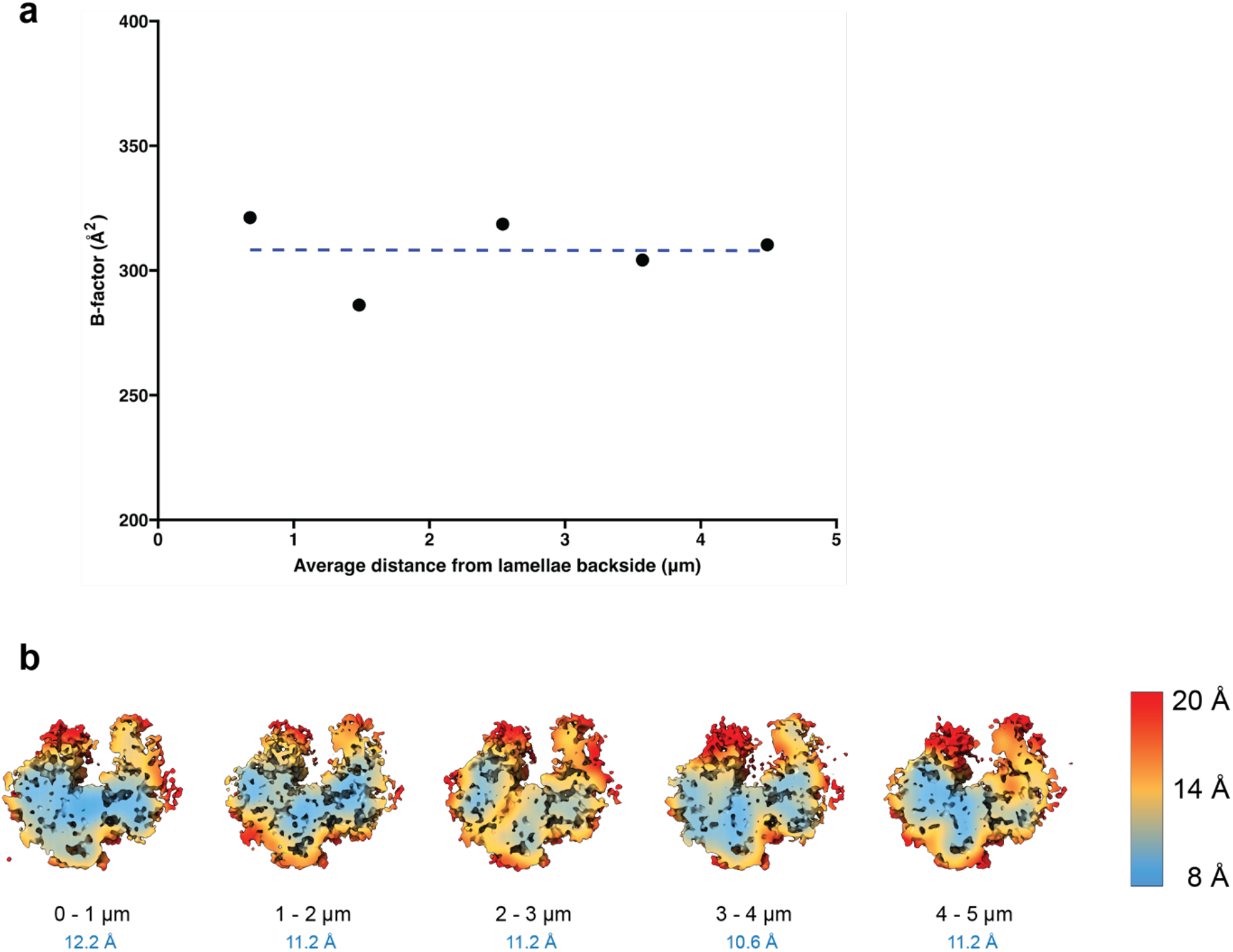
**Effect of distance to backside of lamellae on ribosome STA.** (a) Average distance (µm) from backside of lamellae for particles within 5 µm of backside vs B-factor value per group. No correlation could be found (Spearman correlation: r = -0.3 p-value = 0.68) (b) Local resolution maps for backside damage distance groups, for 800 particles. Global resolution estimates in blue.

**Supplementary Figure 6.**
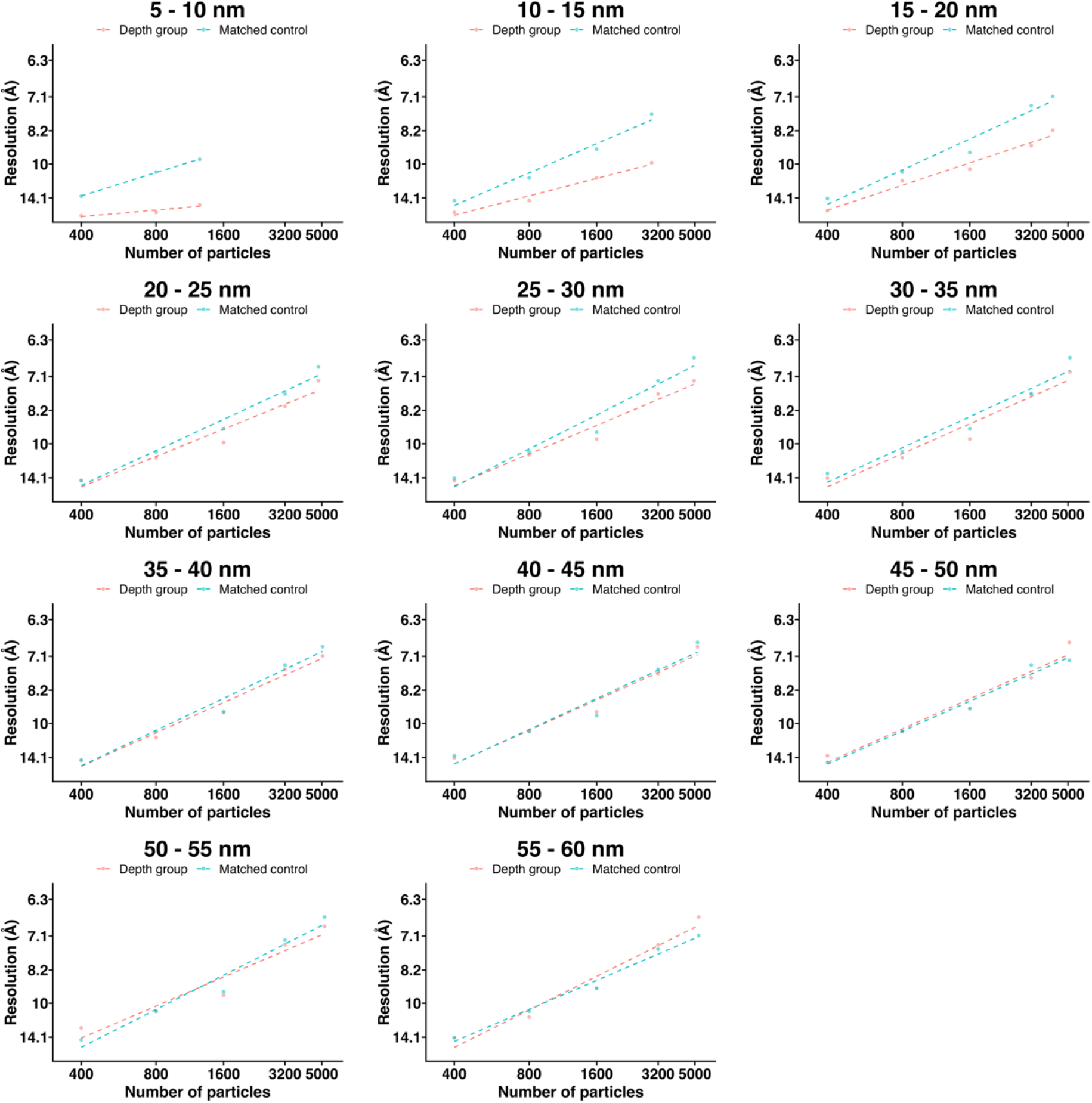
B-factor plots of FIB surface damage groups. B-factor plots for ribosome particles grouped by distance from the PFIB milling surfaces (blue) and their corresponding matched controls of ribosome particles located further from the milling surfaces (orange).

**Supplementary Figure 7.**
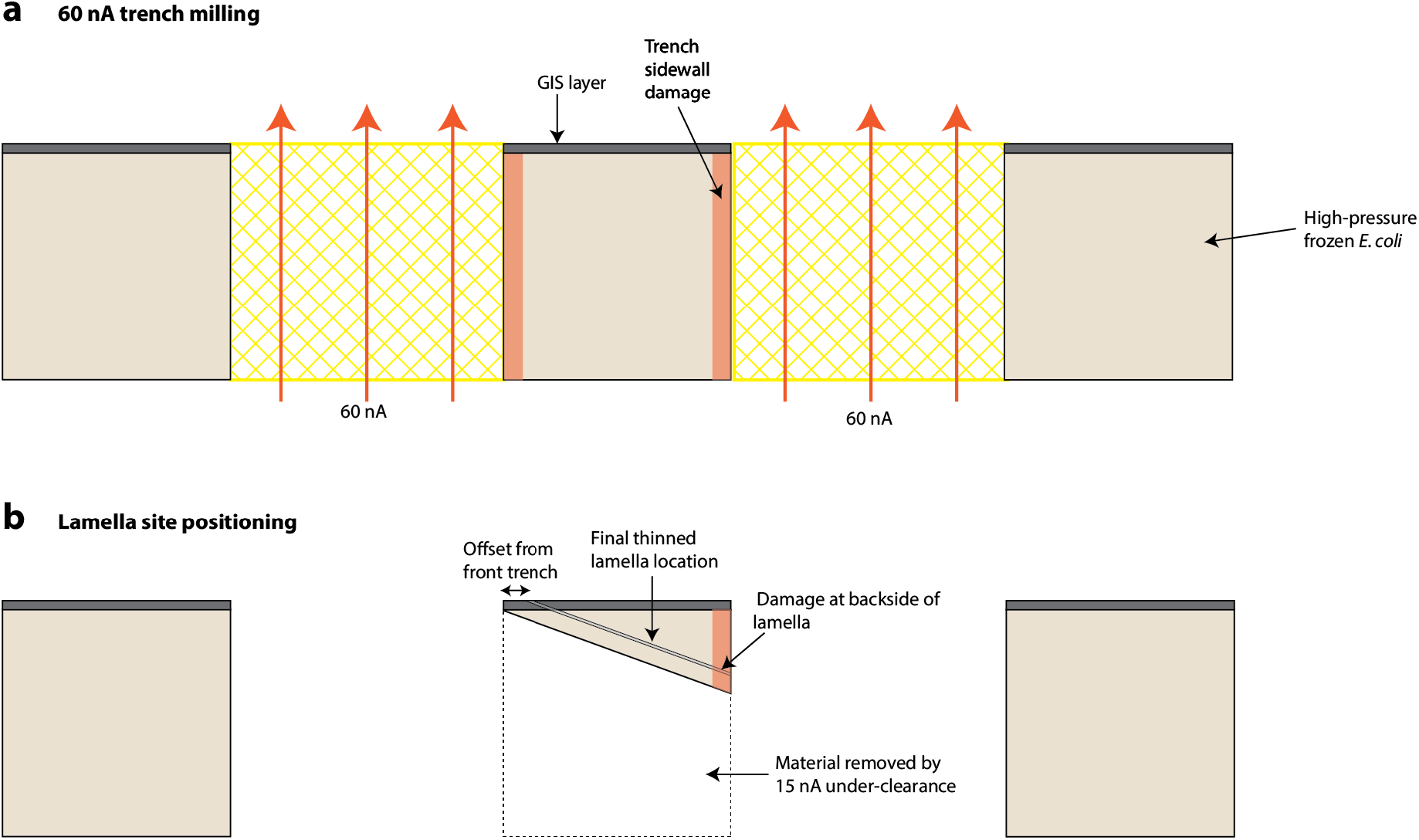
Proposed model for backside damage to lamella. (a) Trenches are milled with 60 nA xenon from the backside of the grid (ablated material indicated in yellow, direction of milling indicated by orange arrows). We speculate this causes a region of damage to the side walls of the trenches (red). **(b)** Following clearance of material under the lamella site at 15 nA, the final lamella site is positioned, with the front edge typically slightly offset from the front trench. The shallow milling angle therefore means that the final lamella includes the damaged sample material (red) on the back side, but not the front.

**Supplementary Figure 8.**
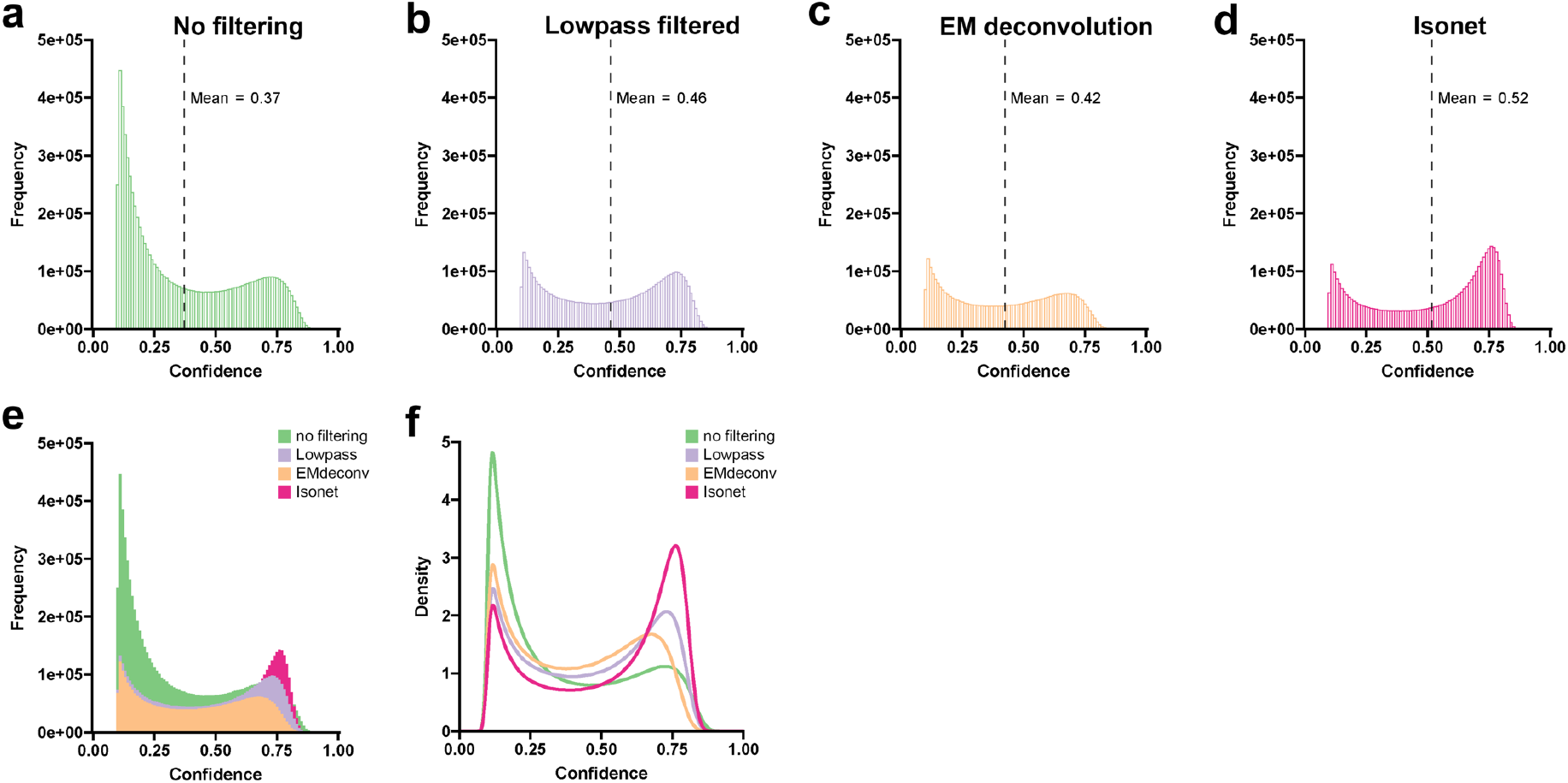
**Effects of filtering on effectiveness of automated ribosome particle picking.** Distribution of crYOLO-assigned confidence scores for machine-learning based picking of ribosomes with crYOLO (Wagner et al., 2019) on filtered or unfiltered tomograms. (a-d) Histograms (bin width 0.01) showing the frequency of the confidence scores during prediction for ribosome positions identified in 2D slices of tomograms that were: SART reconstructed tomograms without additional filtering (a), lowpass filtered (b), EM deconvolution filtered (c) or filtered using machine-learning based missing-wedge compensation as implemented in Isonet (Liu et al., 2022). Values for the mean are shown in the plots and indicated with vertical black dashed lines. A total of 8.01 x 10^6^ 2D ribosome positions were identified on unfiltered tomograms, 4.70 x 10^6^ for lowpass filtered tomograms, 3.65 x 10^6^ for EM deconvolved tomograms and 4.37 x 10^6^ for Isonet filtered tomograms. (e) Overlay of the histograms shown in panel a-d. (f) Density plots for the confidence scores of the same four conditions.

## Supplementary Videos

**Supplementary Video 1.** Removal of Ice contamination from the grid, by imaging with the FIB beam at low magnification.

**Supplementary Video 2.** Tomograms recorded on lamellae with variable local thicknesses, for which a single slice is shown in Figure 2c-e. Scalebar: 100 nm

**Supplementary Video 3.** Movie of consensus STA density map of *E. coli* 70S ribosome surfaces coloured by local resolution. Scalebar: 1 nm. Colour key used is the same as that used in Figure 2. **Supplementary Video 4.** Aligned tilt-series of the striated damage pattern at the back of a lamella shown in Figure 3b. Scalebar: 100 nm.

**Supplementary Video 5.** Reconstructed tomogram of the striated damage pattern at the back of a lamella, from the tilt-series shown in Figure 3b and Supplementary Video 3. Considerable reconstruction artefacts are present at the very back of the lamella, visible as streaks extending from the edges of high-contrast bands of the damaged area. Scalebar: 100 nm.

